# Helarchaeota and Co-occurring Sulfate-Reducing Bacteria in Subseafloor Sediments from the Costa Rica Margin

**DOI:** 10.1101/2021.01.19.427333

**Authors:** Rui Zhao, Jennifer F. Biddle

## Abstract

Deep sediments host many archaeal lineages, including those of the Asgard superphylum that may depend on/require syntrophic partnerships. Our knowledge about sedimentary archaeal diversity and their metabolic pathways and syntrophic partners is still very limited. We present here new genomes of Helarchaeota and co-occurring sulfate-reducing bacteria (SRB) recovered from organic-rich sediments off Costa Rica Margin. Our molecular analyses revealed three new metagenome-assembled genomes (MAGs) affiliating with Helarchaeota, each of which has three variants of the methyl-CoM reductase-like (MCR-like) complex that may enable them to oxidize short-chain alkanes anaerobically. These Helarchaeota have no multi-heme cytochromes (MHCs) but have Group 3b and Group 3c [NiFe] hydrogenases, and formate dehydrogenase, and therefore could transfer the reducing equivalents generated from alkane oxidation to external partners via the transfer of these substances. We also recovered five MAGs of SRB affiliated with the class of Desulfobacteria, two of which showed relative abundances (represented by genome coverages) positively correlated with those of the three Helarchaeota. Genome analysis suggested that these SRB bacteria have the capacity of H_2_ and formate utilizations and may facilitate electron transfers from other organisms by means of these reduced substances. Our findings suggest that Helarchaeota may metabolize synergistically with SRB in marine anoxic sediments, and exert an important influence on the carbon cycle by mitigating the hydrocarbon emission from sediments to the overlying ocean.

## Introduction

Many of the total microbial cells in the marine realm are estimated to be present in marine sediments (Kallmeyer et al., 2012), in which a considerable fraction of the microbial cells are archaea (Lipp et al., 2008; Lloyd et al., 2013a; Hoshino and Inagaki, 2019). Although archaeal communities in oligotrophic and often oxic sediments are dominated by aerobic ammonia-oxidizing Thaumarchaeota (Vuillemin et al., 2019; Zhao et al., 2019; Hiraoka et al., 2020; Hoshino et al., 2020), those in coastal and often organic-rich sediments are more diverse and complex (Durbin and Teske, 2012), and their metabolic activity exerts a critical influence in the carbon and nutrient cycling on the global scale (e.g. (Biddle et al., 2006; Lloyd et al., 2013b; Yu et al., 2018)).

The Asgard superphylum of archaea is of evolutionary importance because its present-day members are thought to share a common ancestor with modern eukaryotes (Spang et al., 2015; Zaremba-Niedzwiedzka et al., 2017). The archaeal phyla of this superphylum, including but not limited to Lokiarchaeota, Thorarchaeota, Odinarchaeota, and Heimdallarchaeota, have been established mainly by metagenome-assembled genomes (Spang et al., 2017; Zaremba-Niedzwiedzka et al., 2017). A recent analysis of available Asgard 16S rRNA gene sequences (Quast et al., 2013) showed that some sequences cannot be resolved to the existing lineages and thus suggested the diversity of Asgard archaea is likely broader than currently recognized (Manoharan et al., 2019). Marine sediments delivered the first cultured Asgard archaeon *Candidatus* Prometheoarchaeum syntrophicum MK-D1 (Imachi et al., 2020) and first identification of the superphylum via metagenomic analysis of DNA (Spang et al., 2015; Zaremba-Niedzwiedzka et al., 2017), so further exploration of marine sediment is promising to recover novel Asgards. This speculation is supported by the recent discovery of Helarchaeota (Seitz et al., 2019), the fifth recognized Asgard phylum, from hydrothermal sediments in the Guaymas Basin (GB).

Genome analysis and laboratory cultures suggest that Asgard archaea in marine sediments will need external partner cells to consume the reducing equivalents (e.g., in the forms of hydrogen and formate) generated during the organic matter degradation of Asgard archaea (Spang et al., 2019; Imachi et al., 2020). Helarchaeota from GB encode the Methyl-CoM reductase-like complex (MCR-like complex) and are proposed to be capable of oxidizing short-chain alkanes to conserve energy (Seitz et al., 2019), using a pathway similar as other thermophilic alkane-oxidizing Euryarchaeota enriched from the same location (Laso-Perez et al., 2016; Chen et al., 2019). Most of these characterized alkane-oxidizing archaea form consortia with sulfate-reducing bacteria (SRB) that use the reducing equivalents released during alkane oxidation for sulfate reduction (Laso-Perez et al., 2016; Chen et al., 2019; Wang et al., 2019), although some alkane-oxidizing archaea can also channel the electrons to their own internal methanogenesis pathway (Laso-Pérez et al., 2019; Wang et al., 2019). As a prevalent SRB in hydrothermal sediments, *Candidatus* Desulfofervidus auxilii (Krukenberg et al., 2016) is the dominant partner bacteria of the characterized alkane-oxidizing archaea in heated marine sediments (Laso-Perez et al., 2016; Chen et al., 2019). However, the potential sinks of the reducing equivalents of Helarchaeota in natural, non-heated sediments still remain elusive.

Scientific drilling off the Costa Rica Margin during International Ocean Drilling Program (IODP) Expedition 334 provided another excellent avenue to explore the diversity of archaea, because sediments from this location were shown to harbor not only extraordinarily high abundances (Martino et al., 2019) but also phylogenetically novel lineages of archaea (Farag et al., 2020b; Farag et al., 2020a). The drilled sites are in the forearc basin of the subduction zone, where a considerable fraction of carbon exchange between Earth’s surface and interior occurs (Barry et al., 2019). Abundant thermogenic alkanes (ethane, propane, and butane) were detected in the deep sediments, but not the surface sediments (Formolo et al., 2011; Expedition 334 Scientists, 2012). This prompted us to perform further metagenomic sequencing and analysis on shallow sediment samples from this expedition, to explore the mechanisms of alkane depletion in this unique setting. Here, we report three new Helarchaeota metagenome-assembled genomes (MAGs) in shallow sediments of Hole U1379B, each of which has three variants of the MCR-like complex and could be the major alkane-oxidizing archaea. These Helarchaeota account for the majority of the mcrA-bearing archaea community, with the rest *mcrA-bearing* archaea affiliated to the family of ANME-1, and may engage in methane metabolism. We also recovered five novel MAGs of sulfate-reducing bacteria in the class of Desulfobacteria, some of which could be the syntrophic partners of Helarchaeota due to the abundance correlation and their matched metabolic capacities of H2 and formate utilization. Our study expands our understanding about the diversity and metabolic functions, and interspecies dependency of Helarchaeota, which could exert a profound influence on the carbon exchange between marine sediments and the overlying seawater.

## Results and Discussion

### Geochemical context

IODP Site U1379 is located on the upper slope of the erosive subduction zone off the Osa Peninsula, Costa Rica. The sediment pile at this site is estimated to be totally 890 meters thick, and characterized by high sedimentation rates of 1.60-10.35 cm/ky (Expedition 334 Scientists, 2012) and low porosities of ~0.6 (Expedition 334 Scientists, 2012). We focused this study on the microbial communities inhabiting 2-9 meters below seafloor (mbsf) of U1379B, including a total of 8 subsampled whole-round cores. Sediments in this depth interval are within the sulfate reduction zone with porewater sulfate concentration decreasing almost linearly with depth (Fig. 1a), while the sulfate-depletion depth was not examined in this study but was noted at ~30 mbsf (Expedition 334 Scientists, 2012). Dissolved Mn concentration in the porewater shows the same decreasing trend as sulfate (Fig. 1b), indicating that Mn^2+^ is mainly consumed rather than produced. The increasing porewater concentrations of ammonium (Fig. 1c) and alkalinity (Fig. 1d) with depth reflect the ongoing degradation of organic matter, likely with sulfate as the most prominent electron acceptor.

**Fig. 1.**
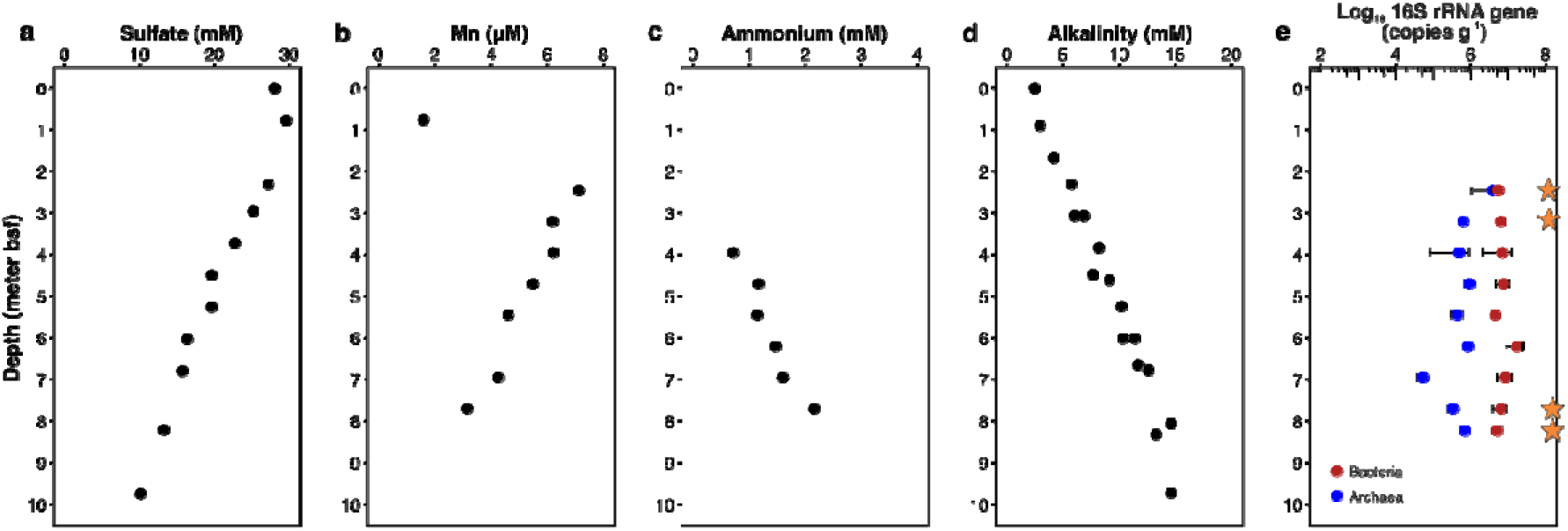
Porewater profiles and microbial abundances in the sediments at IODP Site 1379 on the Costa Rica margin. Porewater profiles were compiled from previous reports/publications of IODP Expedition 334 (Expedition 334 Scientists, 2012; Torres et al., 2014). (a-d) Porewater profiles of sulfate (a), dissolved manganese (b), ammonium (c), and alkalinity (d) in the upper 10 m sediments in Hole U1379. (e) Abundances of archaeal and bacterial 16S rRNA genes quantified using qPCR with domain-specific primers. Stars denote the four sediment horizons where metagenome sequencing data were generated in this study.

### Microbial abundances and overall community structures

We used domain-specific primers (Zhao et al., 2019) to quantify the abundance of archaeal and bacterial 16S rRNA genes in the nine sediment horizons from 2-9 mbsf of Hole U1379B. Both archaeal and bacterial 16S rRNA gene abundances are stable throughout the examined sediments (Fig. 1e). However, archaea are about one order of magnitude lower than bacteria, resulting in the bacterial dominance in most of the studied samples except the shallowest sample (~2 mbsf). A previous study also reported that sediments around ~2 mbsf harbor bacteria and archaea of equal abundances, based on the amplicon and metagenome sequencing data (Martino et al., 2019).

To recover as many archaeal genomes as possible, we selected four sediment horizons (2.45, 3.20, 7.70, and 8.23 mbsf) to subject to shotgun metagenome sequencing, due to the high proportions of archaea in these sediment horizons assessed by qPCR (Fig. 1e). We examined the microbial community structures in these four samples by 16S rRNA gene analyses based on both the un-assembled metagenome reads and the 16S rRNA gene amplicon sequencing data (Fig. 2). Based on the metagenome data, in the archaeal domain, Bathyarchaeota is the major archaeal phylum and accounts for on average 44.7% (37.7-51.2% in individual samples) of the total communities, while Asgard archaea especially Lokiarchaeota were also detected to account for on average 5.6% (2.6-8.7% in individual samples) of the total communities (Fig. 2a). In the bacterial domain, Chloroflexi is the most dominant phylum, which account for on average 19.0% of the total communities in different depths. Other major bacterial phyla include Acidobacteria, Actinobacteria, Elusimicrobia, Planctomycetes and Proteobacteria (mainly Deltaproteobacteria) (Fig. 2a). Similar results were also obtained from the 16S rRNA gene amplicon sequencing and analysis (Fig. 2b). These results are consistent with the previous assessments of microbial community structures of marine sediments off Costa Rica (Martino et al., 2019; Farag et al., 2020b).

**Fig. 2.**
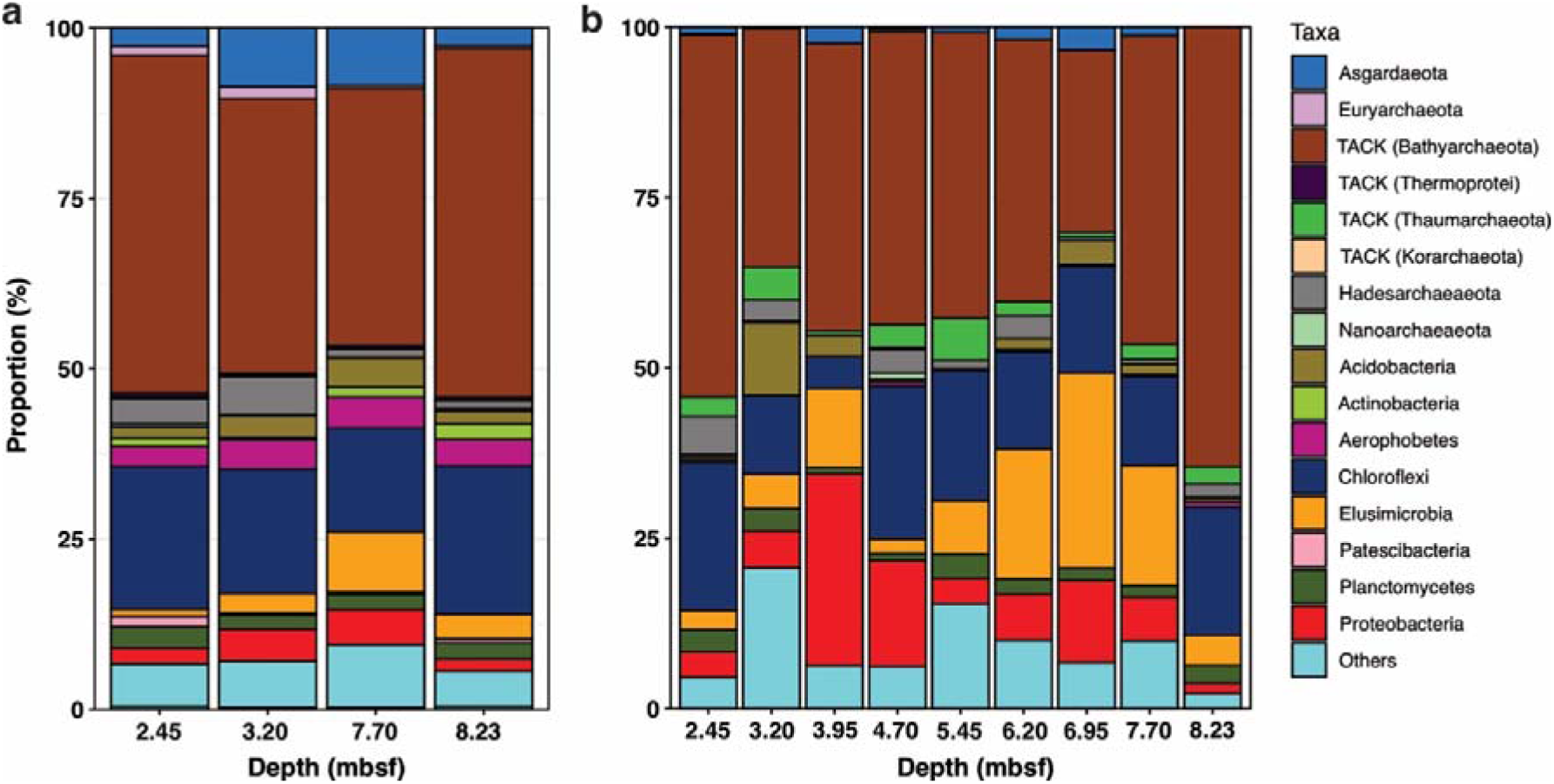
Microbial community structures assessed by 16S rRNA gene reads in raw metagenome sequencing data (a) and amplicon sequencing (b). For (a), unassembled reads were mapped to the SILVA 132 release and classified using phyloFlash (Gruber-Vodicka et al., 2020).

### The presence and diversity of mcrA-bearing microbes

As common products of thermocatalytic degradation of organic matter in subduction zones of high temperature and pressure, C_2_-C_5_ alkanes (ethane, propane, butane, and pentane) were measured in sediments deeper than 50 mbsf (Expedition 334 Scientists, 2012) (Fig. S1). Alkanes in U1379 were generally not detected in the sediments shallower than 50 mbsf and no flux toward the overlying water column can be quantified, suggesting that most of the upward diffusing fluxes of alkanes are consumed in the shallow sediments. To examine the presence and overall diversity of *mcrA-bearing,* potentially alkane-consuming archaea in the CR sediments, we used the GraftM program (Boyd et al., 2018) to detect and classify the *mcrA* gene (encoding the methyl-CoM reductase alpha subunit, the key enzyme of methane/alkane metabolism) in the metagenome datasets. Our results revealed that *mcrA* gene was present in all four metagenome-sequenced sediment samples. Except for 3.20 mbsf, the *mcrA*-containing archaeal communities in all the other three horizons were not dominated by well-known archaea in the Euryarchaeota phylum (Fig. S2 and S3), indicating that uncharacterized archaea may be the dominant methane/alkane metabolizing microbes in the CR sediments, which supports the recently expanded phylogenetic breadth of methane/alkane-metabolizing archaea (Borrel et al., 2019; Hua et al., 2019; Wang et al., 2019).

### Description of the Helarchaeota MAGs

Through metagenome assembly and binning, we obtained 12 archaeal MAGs that could be taxonomically assigned to the Asgard superphylum, three of which are affiliated with the newly proposed Helarchaeota phylum (Seitz et al., 2019), four Lokiarchaeota, and three Heimdallarchaeota (Fig. 3a). The three Helarchaeota MAGs (CR_097, CR_143, and CR_291) recovered from CR have greatly expanded the diversity of this newly proposed phylum, doubling the number of available Helarchaeota genomes (Seitz et al., 2019; Cai et al., 2020). The genome sizes of the three CR Helarchaeota MAGs are similar to SZ_4_Bin10.384 (4.7 Mbp) recovered from coastal sediments (Cai et al., 2020), but are larger than two Helarchaeota genomes recovered from GB (4.7-5.4 Mbp versus 3.54-3.84 Mbp for GB Helarchaeota; Table 1). The CR Helarchaeota MAGs have 4,592 – 4,957 coding sequences distributed on 212 - 589 scaffolds (Table 1). Based on the calculated average amino acid identity (AAI; Table S1), the six available Helarchaeota genomes could be resolved to four families (i.e., with <65% intra-family AAIs; (Konstantinidis et al., 2017)): Hel_GB_A, CR_Bin_097, and CR_Bin_291 belong to a family, while each of the rest three genomes represents a family.

**Fig. 3.**
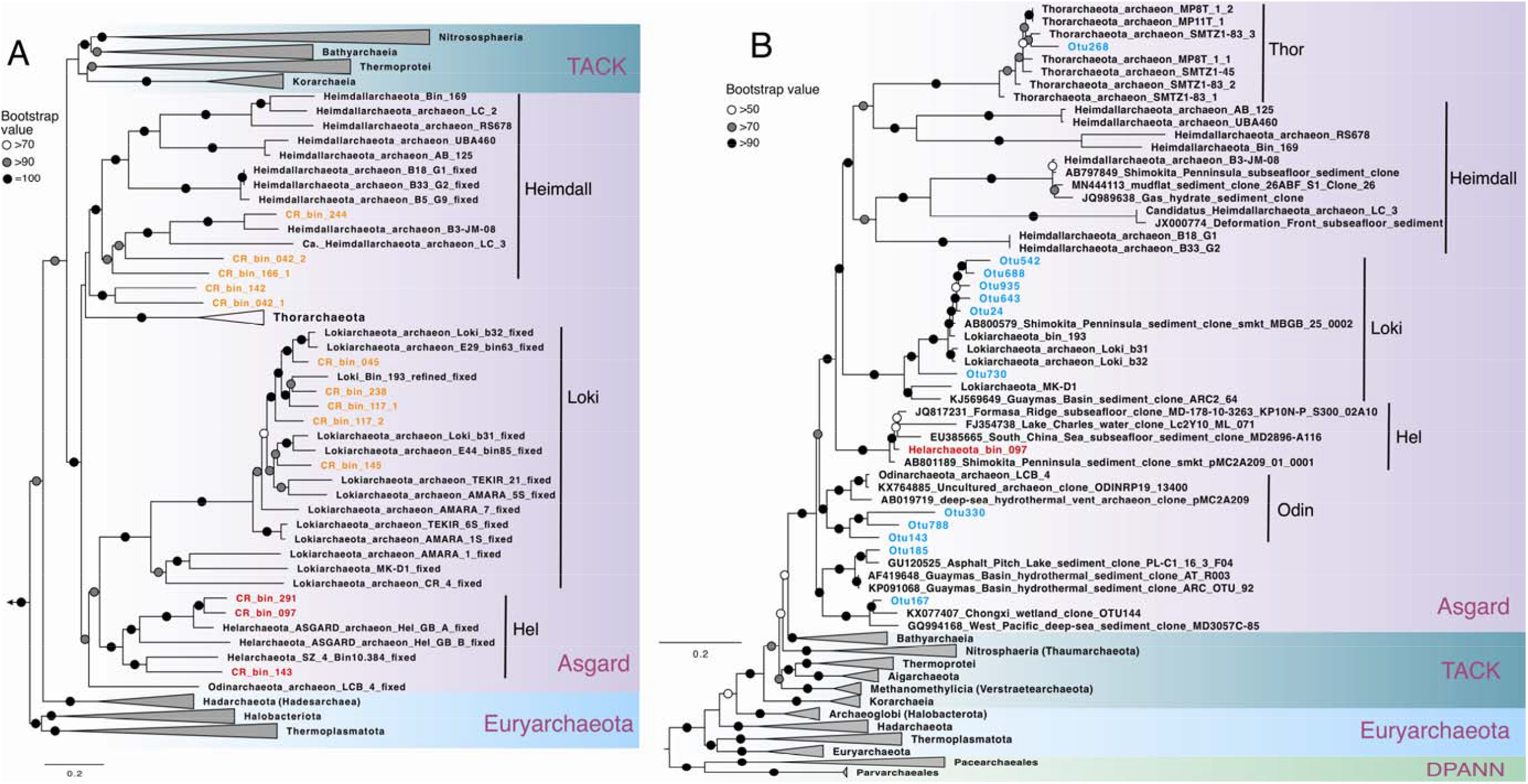
Maximum-likelihood phylogenetic tree of 15 concatenated ribosomal proteins (a) and 16S rRNA gene (b) of archaea. **(a)** This tree was inferred using IQ-TREE v1.6.10 with the LG+R7 model and 1000 ultrafast bootstraps. The Helarchaeota MAGs recovered in this study is highlighted in red. Asgard archaea MAGs other than Helarchaeota recovered in this study are shown in orange. **(a)** This tree was inferred using IQ-TREE v1.6.10 with the SYM+R5 model and 1000 ultrafast bootstraps. The Helarchaeota MAG is highlighted in red, while the short OTUs (287 bp) affiliated with Asgard archaea are shown in blue. Lineages were collapsed at the phylum level if possible, except for those in the Asgard superphylum. The scale bars show estimated sequence substitutions per residue.

**Table 1.**
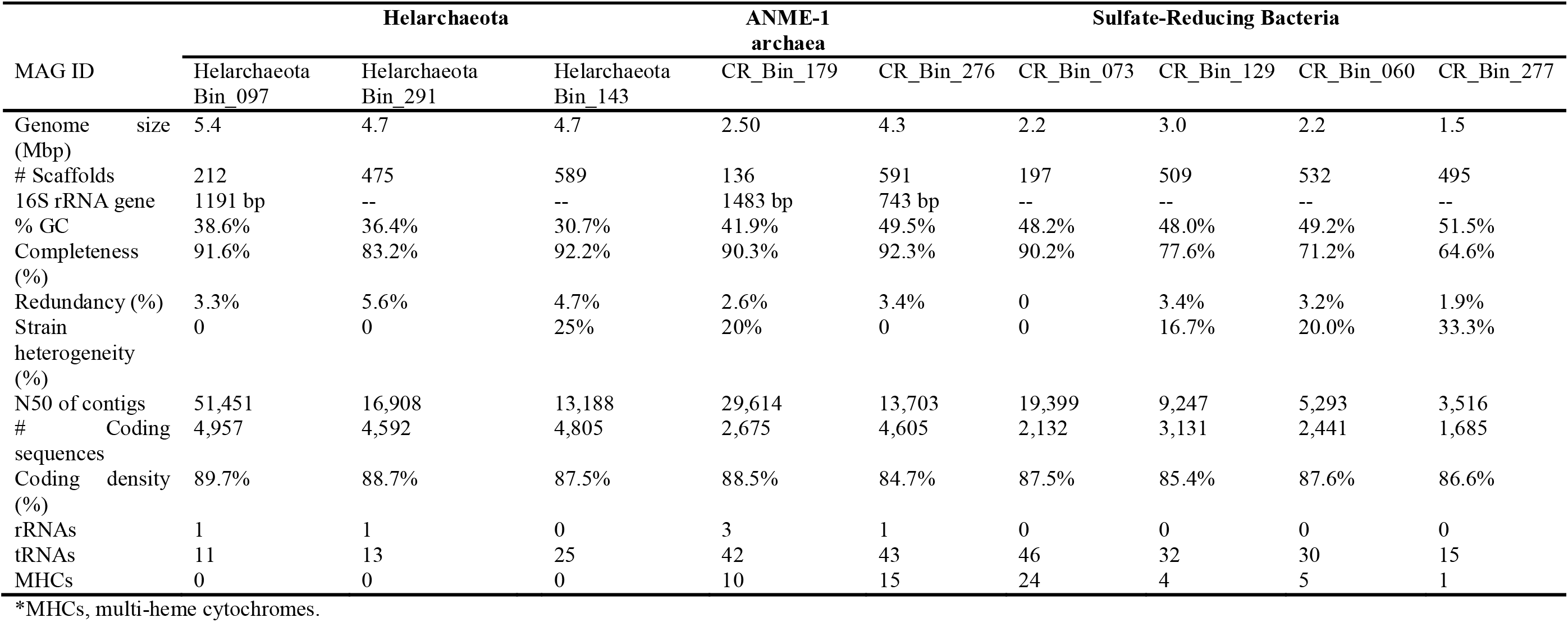
Summary of the Helarchaeota and Deltaproteobacteria MAGs recovered from Hole U1379B

Among the three CR Helarchaeota MAGs, only CR_Bin_097 has a reconstructed partial (1191 bp) 16S rRNA gene sequence. This is notable since none of the three previously reported Helarchaeota MAGs has a 16S rRNA gene sequence (Seitz et al., 2019; Cai et al., 2020). An alignment of the 16S rRNA gene sequence of Helarchaeota CR_Bin_097 and other archaea revealed three insertions of >20 bp in this sequence, and such insertions are also present in other environmental clones of Asgard archaea. This observation suggests that insertions in 16S rRNA genes are common not only in bacteria of candidate phyla radiation (Brown et al., 2015), but also in Asgard archaea. Such insertions unfortunately prevent a perfect match between Helarchaeota CR_Bin_097 and the amplicon operational taxonomy units (OTUs), making it challenging to accurately estimate the distribution of these Helarchaeota in the shallow sediment horizons of Hole U1379. However, based on the read recruiting and genome coverage calculation, the three Helarchaeota seem to have similar distribution patterns and be highest in the depth of 7.70 mbsf sample (Fig. S4). After manually removal of the insertions, the 16S rRNA gene of CR_Bin_097 showed nucleotide identities of >90% with diverse uncultured archaea sequences from marine anoxic sediments and formed a monophyletic clade with them in the phylogenetic tree of archaeal 16S rRNA gene (Fig. 3b), supporting that Helarchaeota is a separate phylum within the Asgard superphylum.

### Alkane oxidation potential of CR Archaea

Similar to the GB Helarchaeota (Seitz et al., 2019), Helarchaeota in CR sediments have the potential of alkane oxidation. Each of the three CR Helarchaeota MAGs has three variants of methyl-CoM reductase-like enzymes (McrABG operon, or MCR) (Fig. 4), whereas the previously reported Helarchaeota MAGs (Seitz et al., 2019; Cai et al., 2020) have only 1 - 2 MCR variants. Phylogenetic analyses of the alpha subunit of methyl-CoM reductase-like enzymes (McrA) revealed that McrA sequences of the six available Helarchaeota MAGs form three clusters (Hel Clusters I, II, and III) within a monophyletic cluster together with homologs of Hadesarchaea, Bathyarchaeota, and *Ca.* Methanoliparia (Fig. 4). These lineages also form a monophyletic clade with McrA homologs of Syntrophoarchaea, *Argoarchaeum ethanivorans*, and *Ethanoperedens thermophilum* (Hahn et al., 2020), all are alkane-oxidizing Euryarchaeota confirmed by laboratory incubations (Laso-Perez et al., 2016; Chen et al., 2019; Wang et al., 2019). Because *mcr*-like genes so far have not been found in any Asgard phylum other than Helarchaeota, *mcr* genes in Helarchaeota, as recently suggested (Hua et al., 2019), may have resulted from horizontal gene transfer events from alkane-oxidizing Euryarchaeota. The three Hel clusters of McrA are divergent, with intra-cluster similarities in the range of 44%-63%. Multiple divergent *mcr*-like genes were also reported in “Ca. Syntrophoarchaeum” genomes, in which the duplicated *mcr*-like genes have evolved to use substrates other than methane, such as butane and propane (Laso-Perez et al., 2016). These CR Helarchaeota MAGs also possess genes associated with the archaeal Wood-Ljungdahl, fatty acids beta-oxidation, and other pathways similar to those found in the characterized alkane-oxidizing genomes (Laso-Perez et al., 2016; Hahn et al., 2020) that have been recently identified (Seitz et al., 2019). Therefore, the gene duplication in Helarchaeota may lead to the neofunctionalization (Rastogi and Liberles, 2005) of the MCR-like complex, and enable them to metabolize short chain alkanes of different lengths anaerobically.

**Fig. 4.**
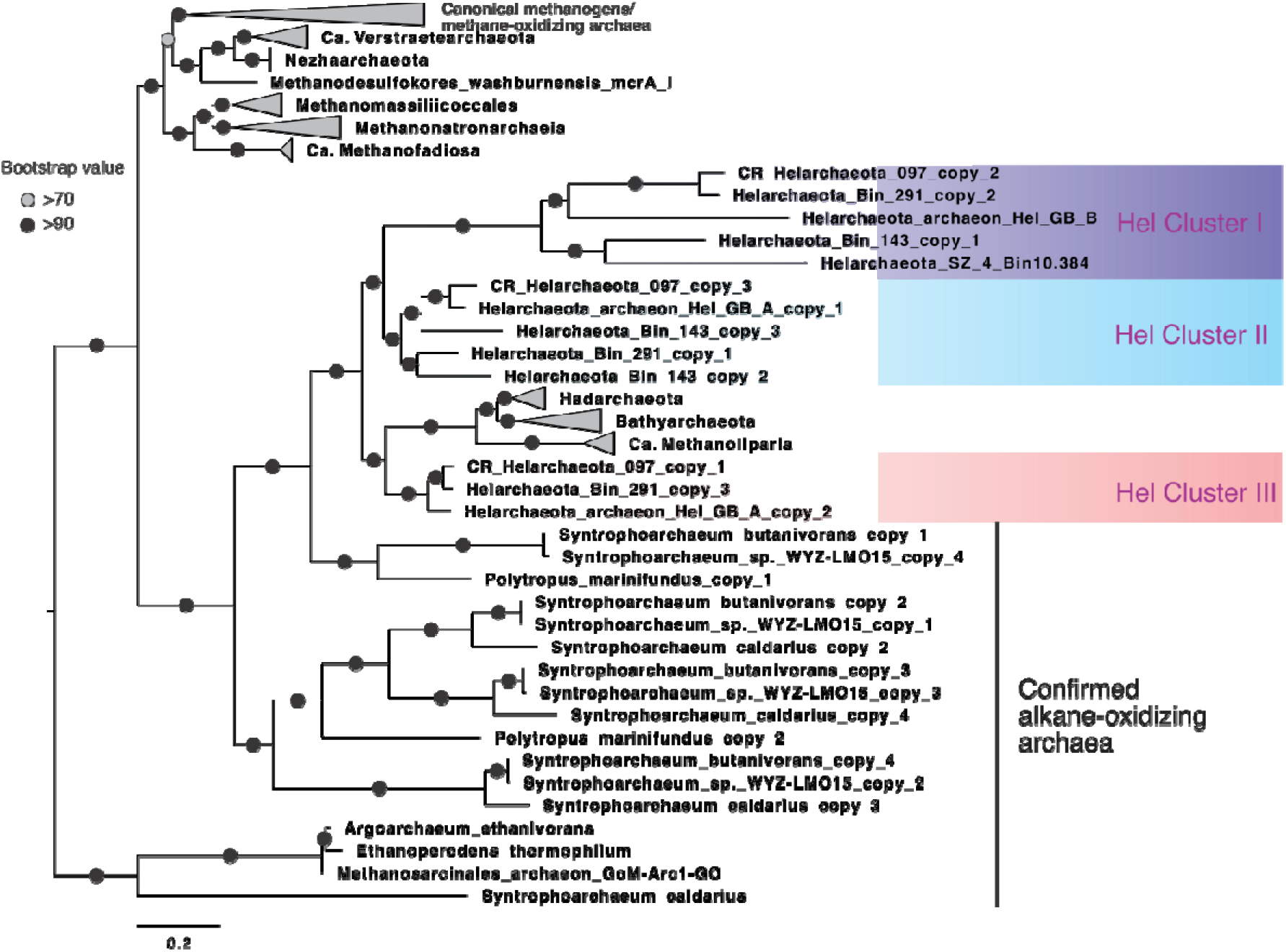
Maximum-likelihood phylogeny of methyl-coenzyme M reductase submit alpha (McrA) of archaea. This tree was inferred using IQ-TREE v1.6.10 with LG+F+I+G4 as the best-fit evolutionary model and 1000 ultrafast bootstraps based on an alignment of 450 amino acid positions of 167 sequences from diverse MCR-containing archaea. The McrA sequences of Helarchaeota were placed into three distinct clusters: Hel Cluster I, II, and III. The scale bar shows estimated sequence substitutions per residue.

Among the MCR-containing MAGs from CR sediments, in addition to the three Helarchaeota MAGs, we also recovered an MAG (CR_Bin_179) affiliated with ANME-1. Based on the phylogenetic analysis of the 15 concatenated ribosomal proteins (Hug et al., 2016), this MAG should represent a new genus within the family of ANME-1 (Fig. S5A), which is also supported by the classification of GTDB-tk (Chaumeil et al., 2020) using the 122 single-copy genes of archaea, and the calculated 69%-76% AAI between this MAG and the others in the family of ANME-1 (Supplementary Table S2). CR_ANME1_Bin_179 has a near full-length 16S rRNA gene sequence (1 487 bp), which shows an identity of 94.9% with that of ANME1_CONS3730B06UFb1 (a MAG recovered from Hydrate Ridge methane seep sediments (Skennerton et al., 2017)) and even lower identities with the other genomes in the family of ANME-1, supporting that this genome should represent a new genus (Konstantinidis et al., 2017). Its genomic close relatives include ANME1_CONS3730B06UFb1 recovered from Hydrate Ridge methane seep sediments, M5.MMPM from Aarhus Bay sediments, and ANME-1-THS recovered from Tibetan hot spring sediments, for all of which methane metabolism pathways have been proposed (Skennerton et al., 2017; Beulig et al., 2019; Borrel et al., 2019). CR_ANME1_Bin_179 may be involved in methane metabolisms, which was supported by the partial MCR operon (constituted by McrB and McrG but the McrA is unfortunately missing) detected in this genome.

To assess the representability of these McrA-bearing genomes in the CR sediments, we performed a phylogenetic analysis of McrA amino acid sequences in the bulk metagenome assemblies of the four CR sediment layers. Our results indicate that all McrA sequences from CR metagenomes were affiliated with either the three Helarchaeota clusters or the ANME-1 cluster (Fig. S3). Because CR_ ANME1_Bin_179 may only be involved in methane but not alkane metabolism, CR Helarchaeota could be responsible for most of the presumed alkane depletion in the surface sediments at U1379.

Alkane-oxidizing archaea need to employ various means to channel the reducing equivalents generated during the oxidation of alkanes to either another internal metabolic pathway (Laso-Pérez et al., 2019) or to the external partner cells (Laso-Perez et al., 2016; Chen et al., 2019; Wang et al., 2019). Helarchaeota MAGs recovered from CR likely have to depend on bacterial partners as external electron sinks because they do not have (1) the internal pathways for the reduction of sulfate or nitrate/nitrite, (2) multi-heme cytochrome (Table 1), or (3) Type IV electric pilins for extracellular electron transfer to solid electron acceptors (e.g. metal oxides). Instead, these CR Helarchaeota MAGs contain Group 3 [NiFe] hydrogenases and formate dehydrogenases that could facilitate the transfer of the reducing equivalents in the forms of H2 and formate. Phylogenetic analysis of the hydrogenase subunit alpha indicate that all Helarchaeot MAGs have one Group 3b [NiFe] hydrogenase and one Group 3c [NiFe] hydrogenase (Fig. 5), except for CR_Bin_291 in which the missing Group 3b hydrogenase was probably unrecovered. Helarchaeota Group 3b hydrogenases formed a monophyletic cluster with homologs of diverse Lokiarchaeota genomes (Spang et al., 2019), but were divergent from other Group 3b lineages (Fig. 5). Similarly, Helarchaeota Group 3c hydrogenases also formed a monophyletic cluster with homologs of diverse Lokiarchaeota and Thorarchaeota genomes (Spang et al., 2019) (Fig. 5). Interestingly, the only hydrogenase in the closed genome of the cultured *Ca.* Prometheoarchaeum syntrophicum MK-D1 (Imachi et al., 2020) is also affiliated with this cluster. *Ca.* Prometheoarchaeum syntrophicum was revealed to depend on partnering cells to consume the reduced equivalents (in the forms of H_2_ (Spang et al., 2019; Imachi et al., 2020). The congruence between the phylogenies of Asgard Group 3 hydrogenases and their genomes suggested that these hydrogenases may be vertically inherited from their ancestor rather than horizontal transferred from other organisms. Even though these Asgard-specific Group 3 [NiFe] hydrogenases are probably not membrane-bound, they could still play an important role in hydrogen production and transfer between organisms, given that the genome of the first cultured Lokiarchaeota has a single Group 3c [NiFe] hydrogenase (Fig. 5).

**Fig. 5.**
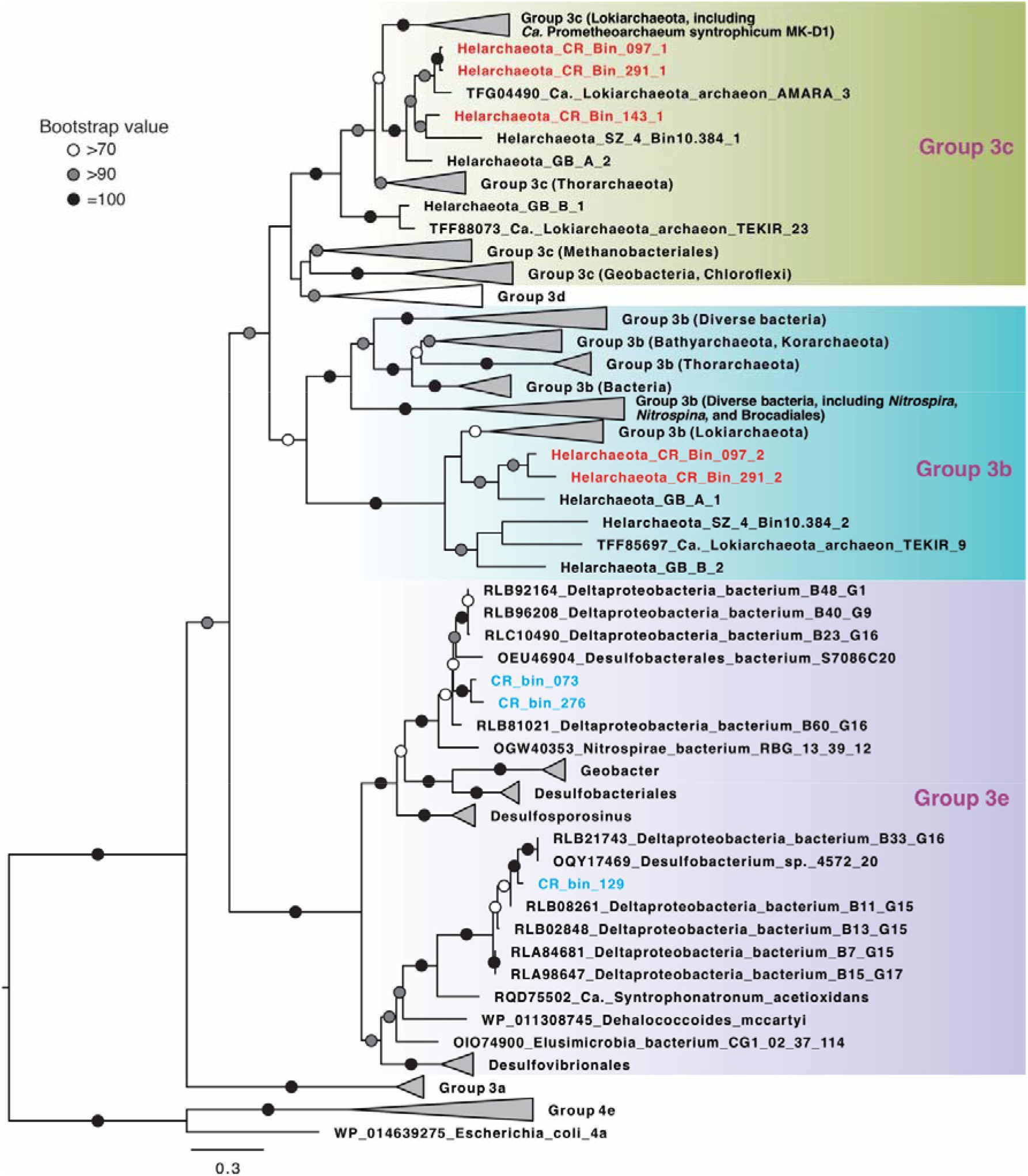
Maximum-likelihood phylogenetic tree of Group 3 [NiFe] hydrogenase alpha subunit. This tree was inferred using IQ-TREE v1.6.10 with LG+R6 as the best-fit evolutionary model and 1000 ultrafast bootstraps based on an alignment of 394 positions of protein sequences from diverse Bacteria and Archaea. Sequences of Helarchaeota recovered from CR are shown in red, while those from SRB MAGs are highlighted in blue. The tree is rooted to Group 4 [NiFe] hydrogenase sequences. The scale bar shows estimated sequence substitutions per residue.

### Co-occurrence of sulfate-reducing bacteria and Helarchaeota

The potential partner bacteria of Helarchaeota have not been identified. From the same metagenome sequencing dataset, we recovered five bacterial MAGs affiliated to the phylum of Desulfobacterota (Fig. 6a and Fig. 7). Four of them have the complete dissimilatory sulfate reduction pathway (constituted by sulfate adenylyltransferase (sat), adenylyl-sulfate reductase (AprAB), and dissimilatory sulfite reductase (DsrAB)), while the incompletion of this pathway in CR_bin_60 could be due to the low genome completion (71% based on bacterial single-copy genes, Table 1). The presence of SRB genomes in these sediments is not surprising, considering that the examined sediments were from the sulfate reduction zone (Fig. 1). Phylogenomic analysis based on the 15 concatenated ribosomal proteins suggested that all the five SRB MAGs from Costa Rica margin sediments are affiliated to the class of Desulfobacteria in the phylum Desulfubacterota, according to the genome-based taxonomy classification frame (Parks et al., 2018), and are different from *Ca.* Desulfofervidus auxilii (Krukenberg et al., 2016), the partner bacterium of thermophilic alkane-oxidizing archaea, which is affiliated to the class of Desulfofervidia (Fig. 6a). CR_bin_073, CR_bin_276, and CR_bin_277 are members of the order of “C00003060” (Fig. 6a), which corresponds to the lineage of SEEP-SRB1c (Skennerton et al., 2017). The close relative genomes in the order of C00003060 are exclusively recovered from hydrocarbon-rich marine sediments, such as cold seep sites on Hydrate Ridge off the Pacific coast (Skennerton et al., 2017), and B60_G16 recovered from GB hydrothermal sediments (Dombrowski et al., 2018). The other two MAGs, CR_bin_129 and CR_bin_060, formed a new lineage within the order of Desulfatiglandales, together with MAGs recovered from hydrocarbon-rich sediments in GB (Dombrowski et al., 2018) and Gulf of Mexico (Dong et al., 2019) (Fig. 6a).

**Fig. 6.**
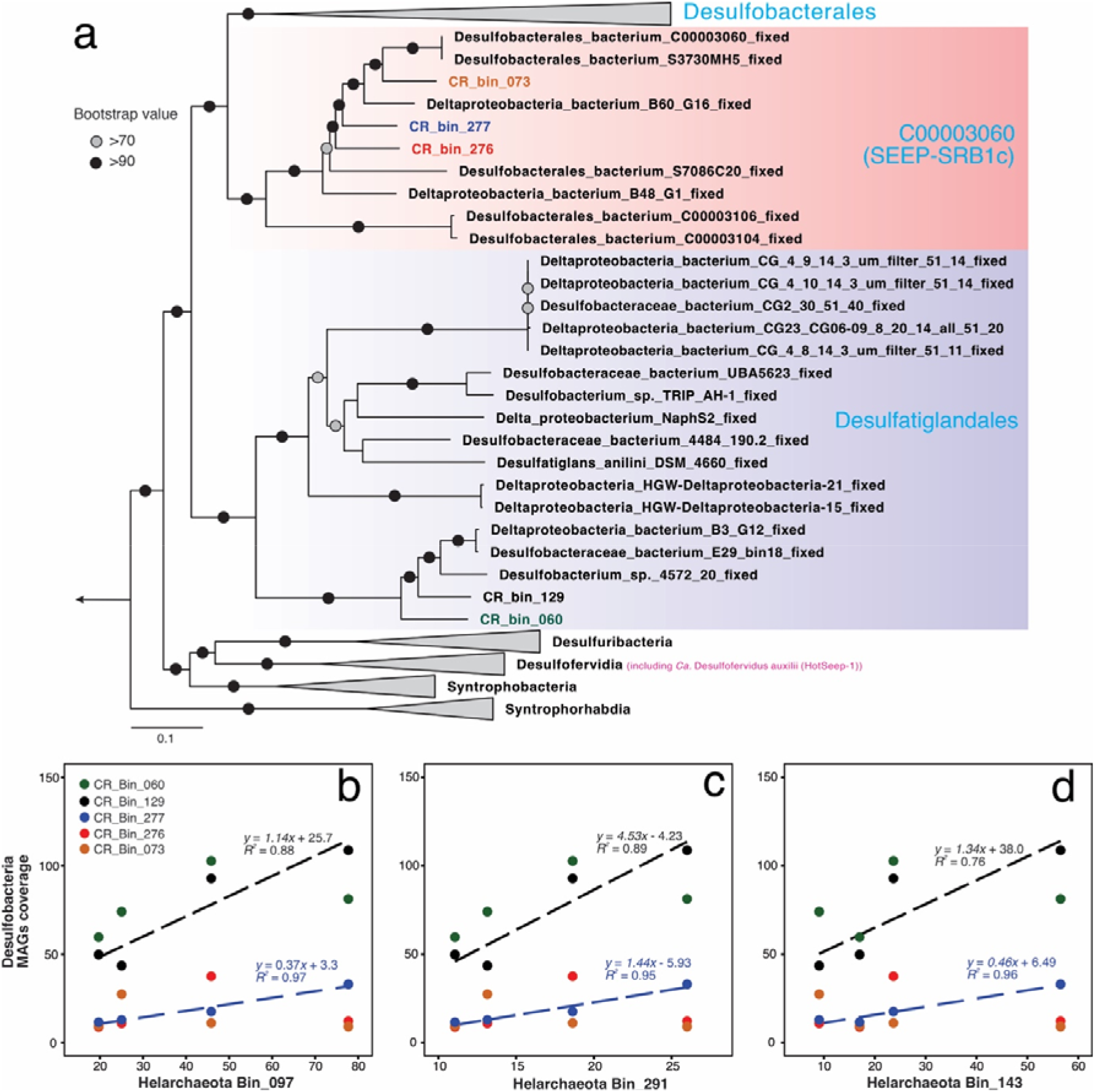
Phylogeny of Desulfobacteria genomes and their genome coverage correlations with Helarchaeota genomes in CR sediments. **(a)** Maximum-likelihood phylogenetic tree of Desulfobacteria genomes recovered from CR sediments. This tree was inferred using IQ-TREE v1.6.10 with LG+F+R6 as the best-fit evolutionary model and 1000 ultrafast bootstraps, based on an alignment of the 15 concatenated ribosomal proteins sequences. This tree is rooted to genomes of the Desulfurmonadota phylum. Desulfobacteria MAGs recovered from Costa Riva margin sediments are color-coded, while the reference genomes are shown in black. Sulfate-reducing bacterial partner of thermophilic alkane-oxidizing archaea, *Candidatus* Desulfofervidus auxilli in the class of Desulfofevidia, is highlighted in purple. The scale bar shows estimated sequence substitutions per residue. **(b-d)** Correlations of the genome coverages between Helarchaeota and Desulfobacteria recovered from CR margin sediments. The blue and black dashed lines show the linear correlations of CR_Bin_277 and CR_Bin_129, respectively, with the three Helarchaeota genomes.

**Fig. 7.**
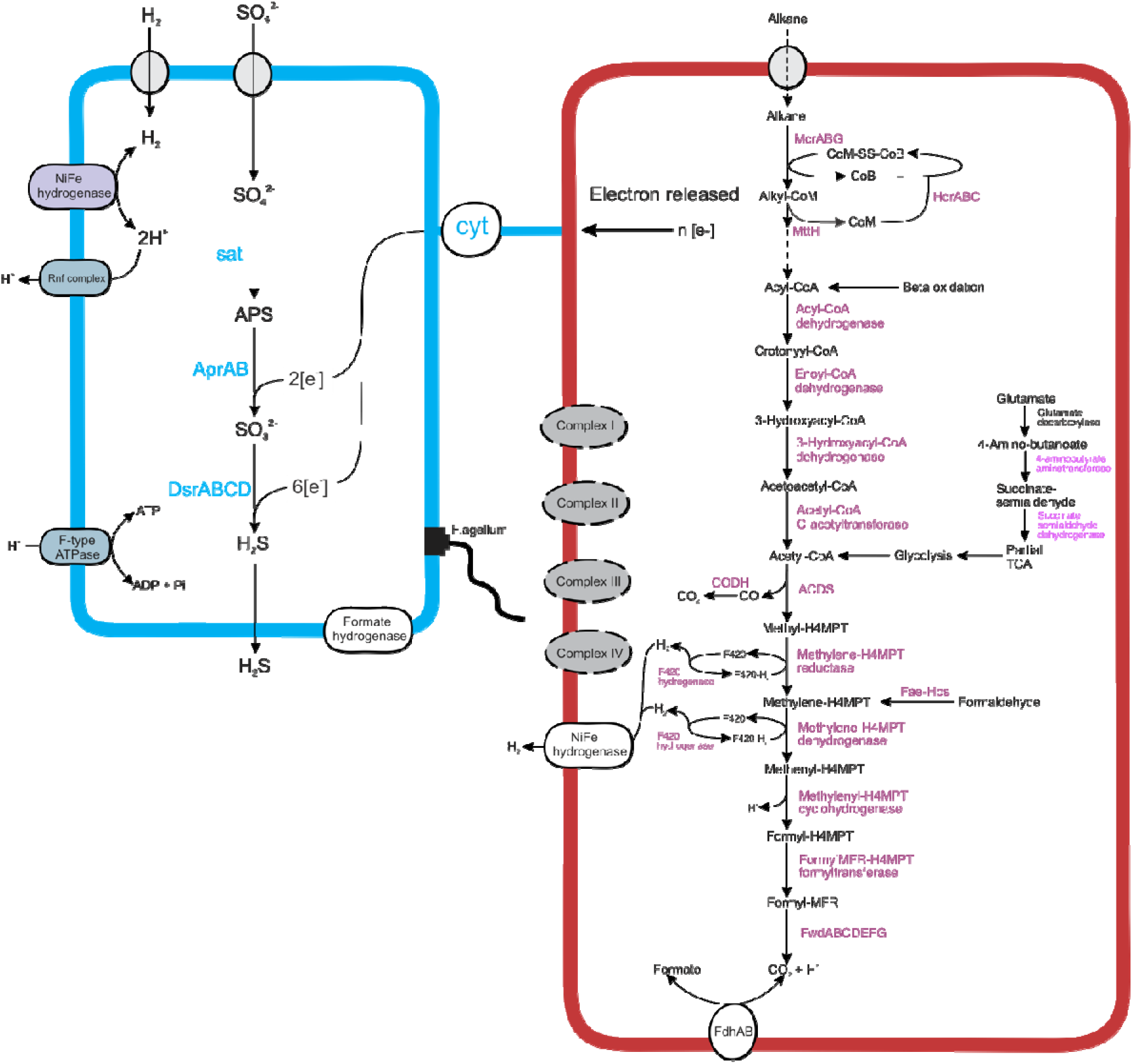
Proposed metabolic scheme and interaction between Helarchaeota (red) and sulfate-reducing bacteria (blue) in Costa Rica Margin sediments. Missing genes/pathways are shown in dashed lines. Cyt, (multi-heme) cytochromes. Absent membrane complexes are shown in grey.

Among the five SRB MAGs, only CR_Bin_276 has an identifiable partial (743 bp) 16S rRNA gene. It shows >99% identity (with two mismatches) with OTU_8 of the amplicon sequencing data. Due to the difficulties to link the Helarchaeota and SRB MAGs with short amplicon OTUs (i.e. Helarchaeota_097 does not match with any OTUs, while only one SRB has a 16S rRNA gene and matches with OTU_8), we use genome coverage as a proxy to represent the relative abundance of MAGs in each of the four metagenome samples. We found that all CR Helarchaeota MAG coverages show positive linear correlations with two of the five SRB MAGs (CR_bin_277 and CR_bin_277) (Fig. 6b-6d), representing the both lineages of SRB recovered from CR. CR_bin_277 and CR_bin_277 co-occur with Helarchaeota in CR sediments.

In addition to the abundance correlations, metabolic pathway analysis of the MAGs also support that the co-occurring Helarchaeota and SRB bacteria in CR sediments may engage in synergy metabolism, in which SRB are capable of utilizing the products of the alkane oxidation activity of Helarchaeota, H2 and formate (Fig. 7). Three of the five SRB MAGs from CR contain a periplasmic hydrogenase affiliated to a novel lineage (Group 3e) in the Group 3 [NiFe] hydrogenases (Fig. 5), which was not well resolved in previous phylogenetic analysis (Greening et al., 2016) probably due to the small number sequences then available. This hydrogenase may enable these SRBs to conserve energy from the oxidation of H_2_, similar as *D. auxilii,* which contains a periplasmic hydrogenase and is capable of growing using H_2_ in the absence of an ANME partner (Krukenberg et al., 2016). In addition, all the five SRB MAGs have two copies of respiratory formate dehydrogenases, indicating that they also have the capacity of oxidizing formate that potential released by the Helarchaeota as an form of reducing equivalents (Fig. 7). These SBR genomes from CR sediments also the full gene set for the synthesis of flagella, which have previously been reported to be an important feature of syntrophy establishment (Shimoyama et al., 2009) and could be important in initiating the contact between SRB and Helarchaeota.

### Sulfate-dependent alkane oxidation is thermodynamically favorable

To explore if the oxidation of short-chain alkanes (ethane, propane, and butane) are thermodynamically favorable when coupled to sulfate reduction, we calculated the Gibbs free energy of these reactions (Supplementary Table S3) under the near *in situ* conditions in the shallow sediments at CR. Our results showed that the Gibbs free energy of the sulfate-dependent alkane oxidation (in the unit of kJ m^-1^ alkane) is feasible over a wide range of alkane concentrations (10^-6^ - 1 μM), and is highest for butane, followed by propane and ethane (Fig. S6). Importantly, these short-chain alkanes can provide more energy than methane per mole of oxidized alkane (Fig. S5), suggesting that these alkanes are more energy-rich than methane and could be more easily consumed by microbes such as Helarchaeota and SRB detected in CR.

Although more data are required to properly assess the ecological roles of Helarchaeota in the environment, all known Helarchaeota so far are found in hydrocarbon-rich marine sediments (Seitz et al., 2019) and seem to be intrinsically related to the oxidation of alkanes. If they have a sulfate-reducing bacterial partner, their syntrophic metabolic reactions, the sulfate-dependent alkane oxidations, are also high energy-yielding processes. Analogous to the role anaerobic methane-oxidizing archaea have on reducing methane emission from marine sediments (Valentine and Reeburgh, 2000), the activities of Helarchaeota and SRB in marine sediments may convert alkane to CO_2_ at the expense of sulfate reduction and, therefore, reduce the emission of alkanes into the overlying water column (Fig. S1).

### Conclusion

This study revealed that the majority of Asgard archaeal cells in sulfate-reducing organic-rich shallow sediments co-occur with SRB in the subduction zone off Costa Rica. Among these are three new Helarchaeota identified as MAGs, each of which have three variants of the MCR operon, and may anaerobically oxidize the steadily available alkane that is thermogenically produced from organic matter degradation in deep sediments. Like other characterized thermophilic alkane-oxidizing archaea, these Helarchaeota may engage in a syntrophic relationship with the co-occurring sulfate-reducing bacteria from the class of Desulfobacteria. These findings expand the diversity of Helarchaeota, and suggest that geological products such as hydrocarbons released from thermogenic processes in subduction zones exemplified by the Costa Rica Margin could play an important role in fueling the catabolism of microbial life in the marine deep biosphere. Experimental approaches, such as laboratory incubation complemented with cell visualization, further microscopic observations and transcriptome sequencing (Laso-Perez et al., 2016) are required to confirm these proposed syntrophic interactions.

## Materials and Methods

### Study sites and sample collection

Sediments used in this study were collected from Hole U1379B at the Costa Rica margin during the IODP Expedition 334. This site is located on the Caribbean plate shelf into the upper slope, with the water depth of this site is 127 m. Sediments in this Hole was drilled using the advanced piston corer system with the core recovery of 100%. Detail of drilling at this site was provided in (Vannucchi et al., 2013). Thorough descriptions of the sediment properties were published elsewhere (Expedition 334 Scientists, 2012; Martino et al., 2019). Whole-round cores of 10-cm long for microbiology analyses were capped upon drilling using blue plastic caps at both ends, and stored in anoxic plastic bags. Microbiology samples were stored at the Gulf Coast Repository of IODP at −80 °C and then were shipped to the University of Delaware and stored in −80°C freezer until further analyses.

We in this study focused on the shallow silty clay sediments of 2-9 mbsf from Hole U1379B, which is 20 m south to Hole U1379C where the most comprehensive geochemical profiles at this station were measured. The geochemical profiles of sediment in this interval of U1379C were reported in previous publications: porewater concentrations of sulfate, ammonium, and alkalinity in Expedition 334 Scientists (2012), and dissolved manganese in (Torres et al., 2014). Procedures for hydrocarbon composition determined by headspace analysis were described in (Riedinger et al., 2019), and the data can be found in the IODP Expedition 334 report (http://publications.iodp.org/proceedings/334/334toc.htm).

### DNA extraction, quantitative PCR, and amplicon analysis

DNA for amplicon sequencing was extracted from 0.5 g of sediment using the PowerSoil DNA extraction kit (MOBIO, CA). A parallel extraction blank without adding sample material in the beginning was also perform to track the potential contamination introduced during the experimental process. DNA for metagenomic sequencing was extracted from about 5 g of sediment (0.5 g was added into each of the 10 lysis tubes) using the PowerSoil DNA extraction kit (MOBIO, CA). For each sample, DNA extracts from the 10 parallel extractions were eluted with 100 μL double-distilled H_2_O into a single 1.5-mL Eppendorf tube and preserved at −20°C for further analyses.

The bacterial and archaeal 16S rRNA genes were quantified using the primer sets Uni341/Uni519 and Uni515F/Arc908r, respectively, combining with the thermal conditions described in (Zhao et al., 2019). For the amplicon preparation, a two-step strategy was employed to prepare the amplicons of the V4 region of the 16S rRNA gene with the primer pair Uni519F/806R and thermal cycling condition described elsewhere (Zhao et al., 2019). Details about the qPCR standard preparation, qPCR reaction chemistry, amplicon sequencing data analysis were presented in the Supplementary Information.

### Metagenome assembly, binning, and genome refinement

Metagenomic libraries (2×150 paired-end) were prepared and sequenced on an Illumina NextSeq 500 sequencing platform (Illumina Inc., San Diego, CA, USA). Quality of the reads and presence of adaptor sequences were checked using FastQC v.0.11.5 (Andrews, 2010) and then trimmed using Trimmomatic v.0.36 (Bolger et al., 2014). Putative 16S rRNA genes in the trimmed metagenome reads were used to assess the microbial community structure using phyloFlash v3.3 beta 2 (Gruber-Vodicka et al., 2020), in which the short reads were identified and classified against the SILVA 132 release.

The quality-controlled paired-end reads were *de novo* assembled into contigs using Megahit v.1.1.2 (Li et al., 2015) with the k-mer length varying from 27 to 117. Contigs longer than 1000 bp were into automatically binned with MaxBin2 v2.2.6 (Wu et al., 2016) using the default parameters. The quality of the obtained genome bins was assessed using CheckM v.1.0.7 (Parks et al., 2015) with the option “lineage_wf”, which uses lineage-specific sets of single-copy genes to estimate completeness and contamination and assigns contamination to strain heterogeneity if amino acid identity is >90%. Genome bins of >50% completeness were manually refined using the *R* package *gbtools* (Seah and Gruber-Vodicka, 2015), based on the GC content, taxonomic assignments, and differential coverages in different samples. To improve the quality of MAGs, metagenome reads of the sample with the highest coverage was detected were mapped onto the MAG contigs using BBmap (Bushnell, 2014), and the aligned reads were re-assembled using SPAdes v.3.12.0 (Bankevich et al., 2012) with the default parameters and minimum contig length of 1000 bp. The resulting scaffolds were visualized and re-binned using *gbtools* (Seah and Gruber-Vodicka, 2015) as described above. The qualities of the resulting MAGs were checked using the CheckM v. 1.0.7 “lineage_wf” command again. Details were presented in the Supplementary Information.

### Overall diversity of *mcrA-bearing* archaea

Reads of *mcrA* in the unassembled metagenome sequencing data were identified and classified using GraftM v0.13.1 (Boyd et al., 2018), with the pre-curated *mcrA* package (including HMM profiles and pre-constructed phylogenetic tree) described in (Singleton et al., 2018) as the reference database (downloaded from https://data.ace.uq.edu.au/public/graftm/7/).

### Genome annotation

Genome annotation was conducted using Prokka v.1.13 (Seemann, 2014), eggNOG (Huerta-Cepas et al., 2016), and BlastKoala (Kanehisa et al., 2016) using the KEGG database. The functional assignments of genes of interest were also confirmed using BLASTp against the NCBI RefSeq database. The metabolic pathways were reconstructed using KEGG Mapper (Kanehisa et al., 2011).

For the multi-heme cytochromes (MHCs) detection, the heme-binding sites of a protein, the CXXCH motif, were counted using the python script “cytochrome_stats.py” described in (Badalamenti et al., 2016) (available at https://github.com/bondlab/scripts) with the amino acid sequences predicted with Prodigal (Hyatt et al., 2010) as the input. Proteins with >3 CXXCH motifs were counted as MHCs following the criteria described elsewhere (Badalamenti et al., 2016; Hernsdorf et al., 2017).

Average nucleotide identify (ANI) between different genomes were calculated using FastANI with default parameters (Jain et al., 2018). Average amino acid identify (AAI) were calculated using CompareM (https://github.com/dparks1134/CompareM) with the “aai_wf” option, in which the protein coding sequences (CDS) predicted by Prodigal (Hyatt et al., 2010) were taken as the input to identify orthologous proteins by all-vs-all reciprocal sequence similarity search with Diamond (Buchfink et al., 2015). The average similarity of the orthologous proteins between the two genomes were taken as the pairwise AAI.

### Phylogenetic analysis

For the phylogenomic analysis of Helarchaeota, representative genomes of all major archaeal lineages described in the GTDB database (Parks et al., 2018) were downloaded and included. For sulfate-reducing bacteria, representative genomes of the candidate bacteria phyla of Desulfobacterota, Desulfobacterota_A, and Desulfuromonadota (phyla per nomenclatures of GTDB; https://gtdb.ecogenomic.org/) were included. The phylogenomic analyses were based on markers consisting of 15 concatenated ribosomal proteins (rpL2, 3, 4, 5, 6, 14, 16, 18, 22, 24 and rpS3, 8, 10, 17, 19) that have been demonstrated to undergo limited lateral gene transfer (Sorek et al., 2007). These selected proteins, among the conservative single-copy ribosomal proteins included in (Campbell et al., 2013), were identified in Anvi’o v.5.5 (Eren et al., 2015) using Hidden Markov Model (HMM) profiles, following the procedure outlined at http://merenlab.org/2017/06/07/phylogenomics/. Sequences were aligned individually using MUSCLE (Edgar, 2004), and alignment gaps were removed using trimAl (Capella-Gutierrez et al., 2009) with the mode of “automated”. Individual alignments of ribosomal proteins were concatenated. The maximum-likelihood phylogenetic tree was reconstructed based on the concatenated alignment using IQ-TREE v1.6.10 (Nguyen et al., 2015) with LG+F+R6 as the best-fit evolutionary model selected by ModelFinder (Kalyaanamoorthy et al., 2017) and 1000 ultrafast bootstraps using UFBoot2 (Hoang et al., 2018).

A maximum likelihood phylogenetic tree based on 16S rRNA genes was also constructed based on an alignment of 16S rRNA gene sequences of the genomes that included in the above-mentioned phylogenomic analysis. In addition, amplicon OTUs (287 bp) of Asgard archaea and their close relatives (environmental clones of >1300 bp) identified via BLASTn (Altschul et al., 1997) in the NCBI database. Sequences were aligned using MAFFT-LINSi (Katoh and Standley, 2013), and putative insertions were manually trimmed in Unipro UGENE (Okonechnikov et al., 2012). The maximum-likelihood phylogenetic tree was inferred based on the insertion-free alignment using IQ-TREE v1.6.10 (Nguyen et al., 2015) with SYM+R5 as the best-fit evolutionary model determined by ModelFinder (Kalyaanamoorthy et al., 2017) and 1,000 ultrafast bootstrap replicates using UFBoot2 (Hoang et al., 2018).

For the phylogeny of McrA, in addition of Helarchaeota, the genomes of known MCR-bearing archaea genomes were downloaded from NCBI, annotated using Prokka v1.13 (Seemann, 2014), and the McrA amino acid sequences were extracted. Phylogenetic analysis was also performed for the McrA sequences in the bulk assemblies of the four CR sediment horizons, in which the McrA sequences were extracted from the Prokka annotation outputs. In both analyses, all sequences were aligned using MAFFT-LINSi (Katoh and Standley, 2013), trimmed using trimAl (Capella-Gutierrez et al., 2009) with the mode of “automated”, and the maximum likelihood phylogenetic tree was inferred using IQ-TREE v1.6.10 (Nguyen et al., 2015) with LG+F+R6 as the best-fit evolutionary model and 1,000 fast bootstrap replicates.

For the phylogeny of [NiFe] hydrogenases, reference sequences were mainly extracted from (Carnevali et al., 2019) and (Kessler et al., 2019). [NiFe] hydrogenases of the genomes reported in this study were extracted from the Prokka annotations, and used as queries in BLASTp search (Altschul et al., 1997) in the NCBI database to identify their close relatives. All retrieved sequences were aligned using MAFFT-LINSi (Katoh and Standley, 2013), trimmed using trimAl (Capella-Gutierrez et al., 2009) with the mode of “automated”, and the phylogenetic tree was reconstructed using IQ-TREE v1.6.10 (Nguyen et al., 2015), with the procedure described above.

### Genome coverage calculation and the linear correlation

As an proxy of the relative abundance, genome coverages of MAGs in each of the four metagenome-sequenced sediment depths were determined by recruiting reads from the individual metagenome datasets using BBmap (Bushnell, 2014) with the read identity threshold of 98%. Relative abundances of the Helarchaeota MAGs in the total communities were also calculated as the product of the average coverage and genome size, divided by the total trimmed reads. Linear correlations between the five Desulfobacteria MAGs and the three Helarchaeota MAGs were determined in *R* (R Development Core Team, 2011), using Pearson correlation test.

### Thermodynamic calculation

We assessed the feasibility of alkane oxidation processes by calculating the Gibbs free energy for reactions of sulfate-dependent oxidation of ethane, propane, butane as well as methane in the shallow sediments (<10 mbsf) (See Table S3 for the chemical equations). Details were presented in the Supplementary Information.

### Data availability

All sequencing data used in this study are available in NCBI Short Reads Archive under the project number PRJNA599172. In particular, the amplicon sequencing data can be accessed through the BioSample number SAMN13740702 - SAMN13740710. The raw metagenomic sequencing data are available in NCBI under the BioSample numbers SAMN13740741-SAMN13740744. The MAGs described in this study are available in NCBI with the accession numbers JABXJT000000000 - JABXKA000000000.

## Acknowledgements

This study uses samples and data provided by the IODP. We are grateful to all scientists and crew members of IODP Expedition 334, whose efforts made the samples available. We acknowledge computational support from the University of Delaware Center for Bioinformatics and Computational Biology Core Facility and the use of the BIOMIX computing cluster was made possible through funding from Delaware INBRE (NIGMS P20GM103446), the State of Delaware, and the Delaware Biotechnology Institute. This work was funded by the WM Keck Foundation (to J.F.B.).

