## Supplementaty Information for "Helarchaeota and Co-occurring Sulfate-Reducing Bacteria in Subseafloor Sediments from the Costa Rica Margin"

**Supplementary Text**

**Materials and Methods**

***16S rRNA gene amplicon sequencing preparation and sequence analysis***

A two-step strategy was employed to prepare the amplicons of the V4 region of the 16S rRNA gene with the primer pair Uni519F/806R and thermal cycling condition described elsewhere [1]. In the first-round PCR, to obtain similar PCR products across, 1 µl DNA template and 34 PCR cycles were applied for 2H-1, while 3 µl DNA template and 37 cycles were applied for the remaining samples. Duplicate PCR reactions were pooled and purified using GenElute DNA purification kit (Sigma, USA). In the second round PCR, barcodes were attached in the primer with the same nucleotide sequences as the first-round ones. Concentrations of PCR products after purification were measured with Qubit (Thermo Fisher Scientific, USA), and equal amounts of DNA from all samples were pooled. The amplicons were sequenced (2×150 bp paired-ends) on an Illumina NextSeq 500 sequencing platform (Illumina Inc., San Diego, CA, USA).

Sequencing reads were quality filtered and trimmed using the USEARCH v.11 [2] and chimera were detected and removed using UCHIME [3]. Paired-end reads were merged (287 bp) using USEARCH [2]. Trimmed reads were clustered into operational taxonomy units (OTUs) at >97% nucleotide sequence identity using UPARSE [2]. Samples were subsampled to 17,000 reads for each sediment horizon with the –otutab-norm command in USEARCH v.11 [2]. The taxonomic classification of OTUs was performed using the lowest common ancestor algorithm implemented in the CREST package [4] with the SilvaMod132 database as reference.

***Quantitative PCR***

The bacterial and archaeal 16S rRNA genes were quantified using the primer sets Uni341/Uni519 and Uni515F/Arc908r combining with the thermal conditions described in [1]. All qPCR reactions were run in triplicate and each reaction mixture contained 1× QuantiTech SybrGreen PCR master mixture (QIAgen, Germany), 0.5 μM forward and reverse primer and 1 μl of DNA template in a final volume of 20 μL. The standard for each gene was linear genomic DNA from *E. coli* (bacterial 16S rRNA gene) or *Methanosarcina mazei* (archaeal 16S rRNA gene). For each gene, the DNA concentration of the standard was measured using Qubit and a DNA abundance gradient of 10-10^8^ copies µL^-1^ was prepared by 10x serial dilution. Gene abundances were normalized to copies per g sediment.

***16S rRNA gene PCR amplification and sequence analysis***

A two-step strategy was employed to prepare the amplicons of the V4 region of the 16S rRNA gene with the primer pair Uni519F/806R and thermal cycling condition described elsewhere [1]. In the first-round PCR, to obtain similar PCR products across, 1 µl DNA template and 34 PCR cycles were applied for 2H-1, while 3 µl DNA template and 37 cycles were applied for the remaining samples. Duplicate PCR reactions were pooled and purified using GenElute DNA purification kit (Sigma, USA). In the second-round PCR, barcodes were attached in the primer with the same nucleotide sequences as the first-round ones. Concentrations of PCR products after purification were measured with Qubit (Thermo Fisher Scientific, USA), and equal amounts of DNA from all samples were pooled. The amplicons were sequenced (2×150 bp paired-ends) on an Illumina NextSeq 500 sequencing platform (Illumina Inc., San Diego, CA, USA).

***Assembly, binning, and genome refinement***

Metagenomic libraries were prepared and sequenced on an Illumina NextSeq sequencing platform (Illumina Inc., San Diego, CA, USA) at the Genome Sequencing & Phylotype Center in the University of Delaware. Quality of the reads and presence of adaptor sequences were checked using FastQC v.0.11.5 [5]. Then the sequencing data were processed with Trimmomatic v.0.36 [6] to trim read-through adapters (ILLUMINACLIP:TruSeq2-PE.fasta:2:30:10), trim low quality base calls at the starts and ends of reads (LEADING:3, TRAILLING:3), remove reads that had average phred score lower than 25 in a sliding window of 10 bp (SLIDINWINDOW:10:25), and finally remove reads shorter than 100 bp (MINLEN:100). The overall quality of processed reads was evaluated in a final check with FastQC v.0.11.5 [5], to ensure only high-quality reads were used in the downstream analysis.

The quality-controlled paired-end reads were *de novo* assembled into contigs using Megahit v.1.1.2 [7] with the k-mer length varying from 27 to 117. Contigs larger than 1000 bp were into automatically binned with MaxBin2 v2.2.6 [8] using the default parameters. The quality of the obtained genome bins was assessed using the option “lineage_wf” of CheckM v.1.0.7 [9], which uses lineage-specific sets of single-copy genes to estimate completeness and contamination and assigns contamination to strain heterogeneity if amino acid identity is >90%. Genome bins of >50% completeness were manually refined using the R package *gbtools* [10], based on the GC content, taxonomic assignments, and differential coverages in different samples. Coverages of contigs in each sample were determined by mapping trimmed reads onto the contigs using BBMap v.37.61 [11]. Taxonomy of contigs were assigned according to the taxonomy of the single-copy marker genes in contigs identified using a script modified from blobology [12] and classified by BLASTn [13]. SSU rRNA sequences in contigs were identified using Barrnap [14], and classified using VSEARCH [15] with the SILVA 132 release [16] as the reference. To improve the quality of MAGs, metagenome reads of the sample in which the highest coverage was detected were mapped onto the MAG contigs using BBmap [11], and the aligned reads were re-assembled using SPAdes v.3.12.0 [17] with the default parameters and minimum contig length of 1000 bp. The resulting scaffolds were visualized and re-binned using gbtools [10] as described above. The qualities of the resulting MAGs were checked using the CheckM v.1.0.7 “lineage_wf” command again.

**Thermodynamic calculation**

We calculated the Gibbs free energy for reactions of sulfate-dependent oxidation of ethane, propane, butane as well as methane in the shallow sediments (<10 mbsf) (See Table S3 for the chemical equations). The standard Gibbs energy of formation was calculated using the *R* package *CHNOSZ* [18], as corrected for the near in situ temperature (average bottom seawater temperature of 22.6^o^C [19] and pressure (12.5 air pressure, calculated from the water depth of 125 m of this site). In the Gibbs free energy calculation, we considered a wide range of alkane concentrations (10^-9^ – 10^-3^ mM). Sulfate concentration was assumed to 15 mM. Hydrogen sulfide (H_2_S) was not detected in the porewater, but was assumed to be 0.01 µM in our calculation considering that it is generally not detectable in sulfate-bearing sediment of this area [20]. Concentrations of proton and bicarbonate were calculated from pH (~7.8) and DIC (2.5 mM) measurements.

**Supplementary Figures and Tables**

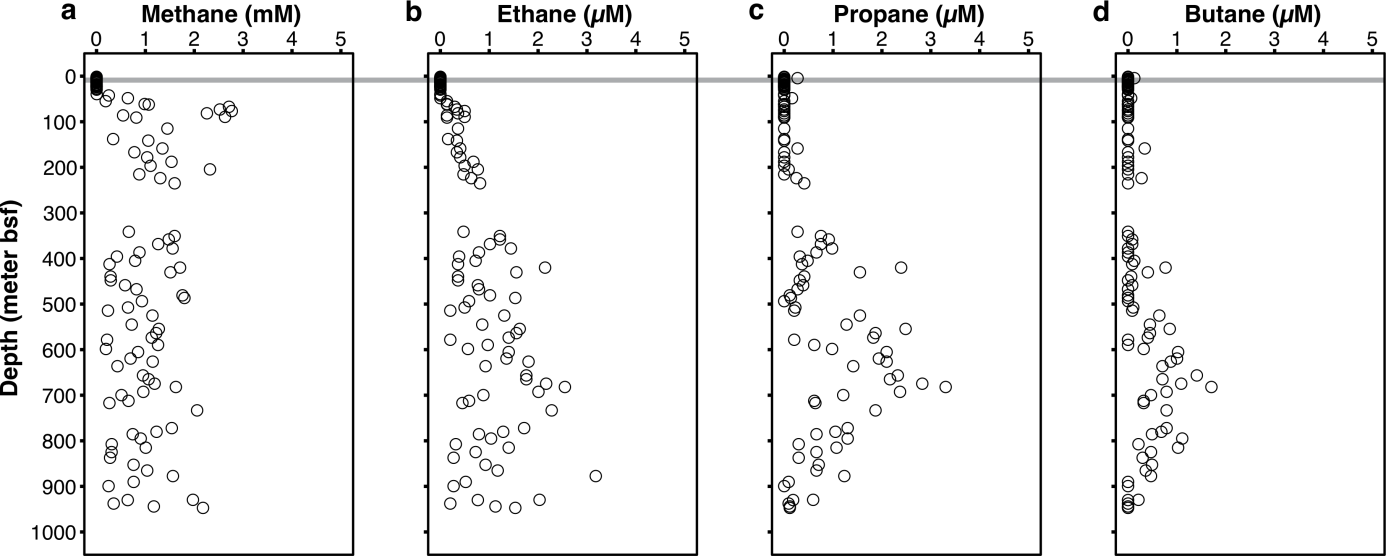

**Fig. S1. Sediment alkane concentrations at IODP Site 1379 off the Costa Rica margin.** Geochemical data were compiled from IODP Expedition 334 report [19]. The thin grey band indicates the sediment interval (2.0-8.2 meters below seafloor) where microbial communities were analyzed in this study.

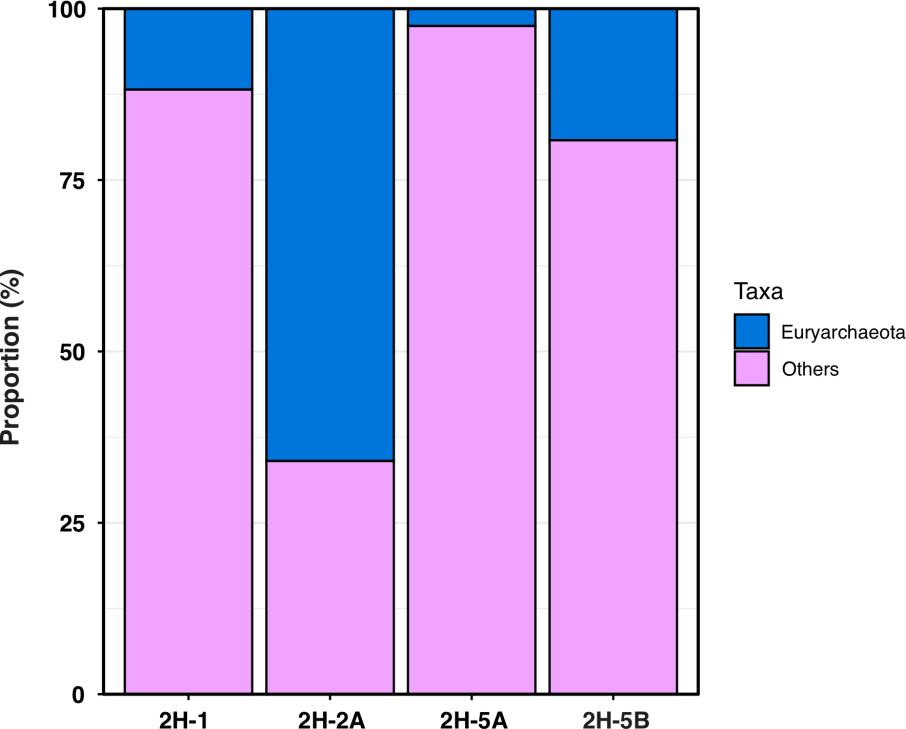

**Fig. S2. Community structure of *mcrA*-bearing microbes in the four selected horizons of IODP Site 1379.** The community structures were assessed based on the un-assembled metagenome reads using GraftM with the curated package of *mcrA* available at <https://data.ace.uq.edu.au/public/graftm/7/>.

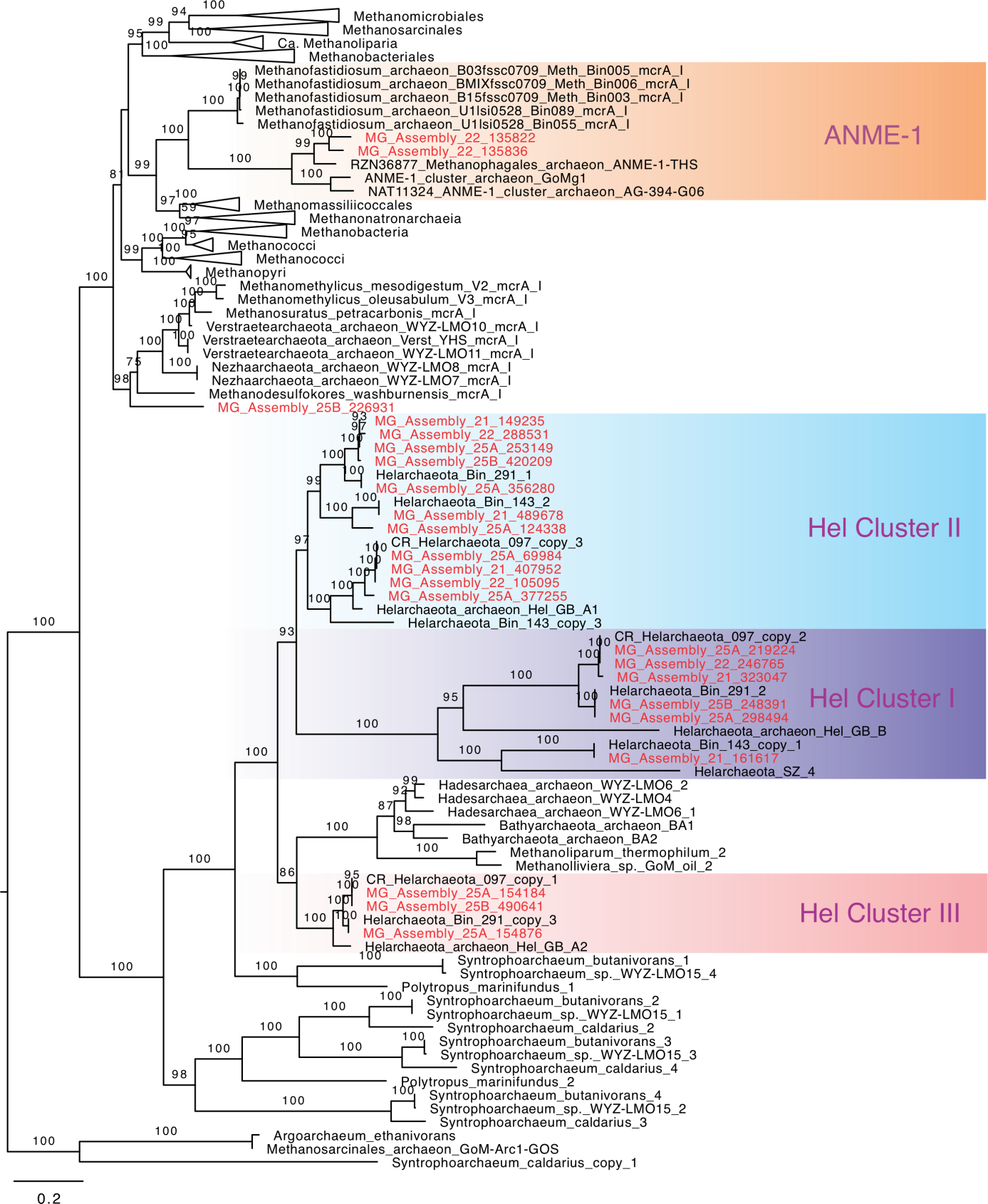

**Fig. S3. Maximum-likelihood phylogenetic tree of methyl-coenzyme M reductase-like complex alpha subunit (McrA).** The tree was reconstructred using IQ-TREE with LG+R6 as the best-fit evolutionary model and 1,000 ultrafast bootstrap iterations. Sequences from contigs of the four metagenome assemblies are shown in red.

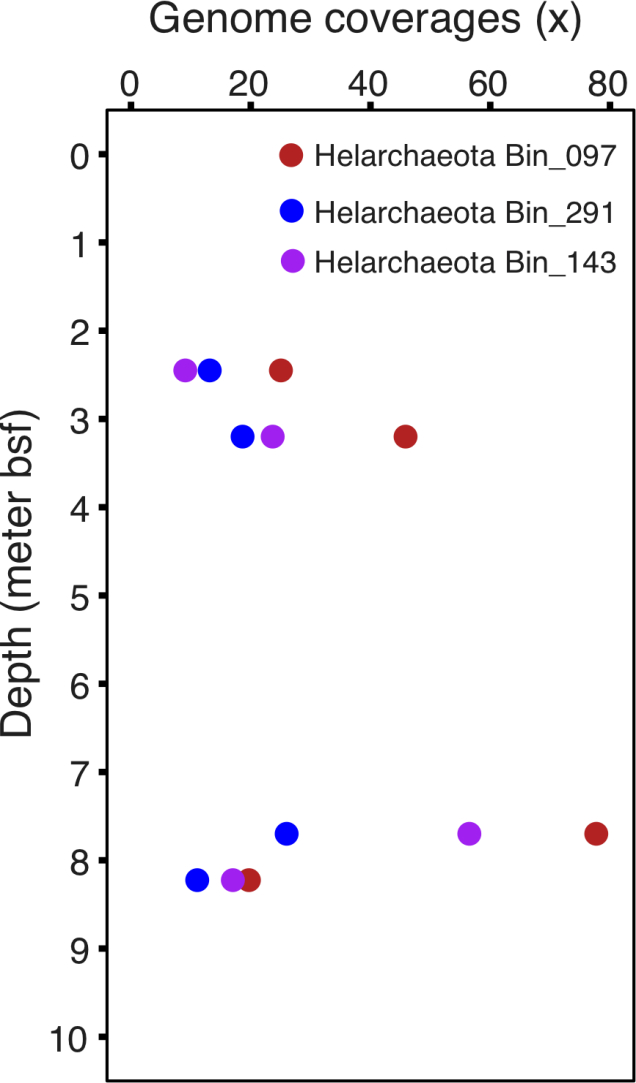

**Fig. S4. Genome coverages of Helarchaeota genomes in different sediment horizons of U1379B.** Genome coverages were determined by reads recruiting.

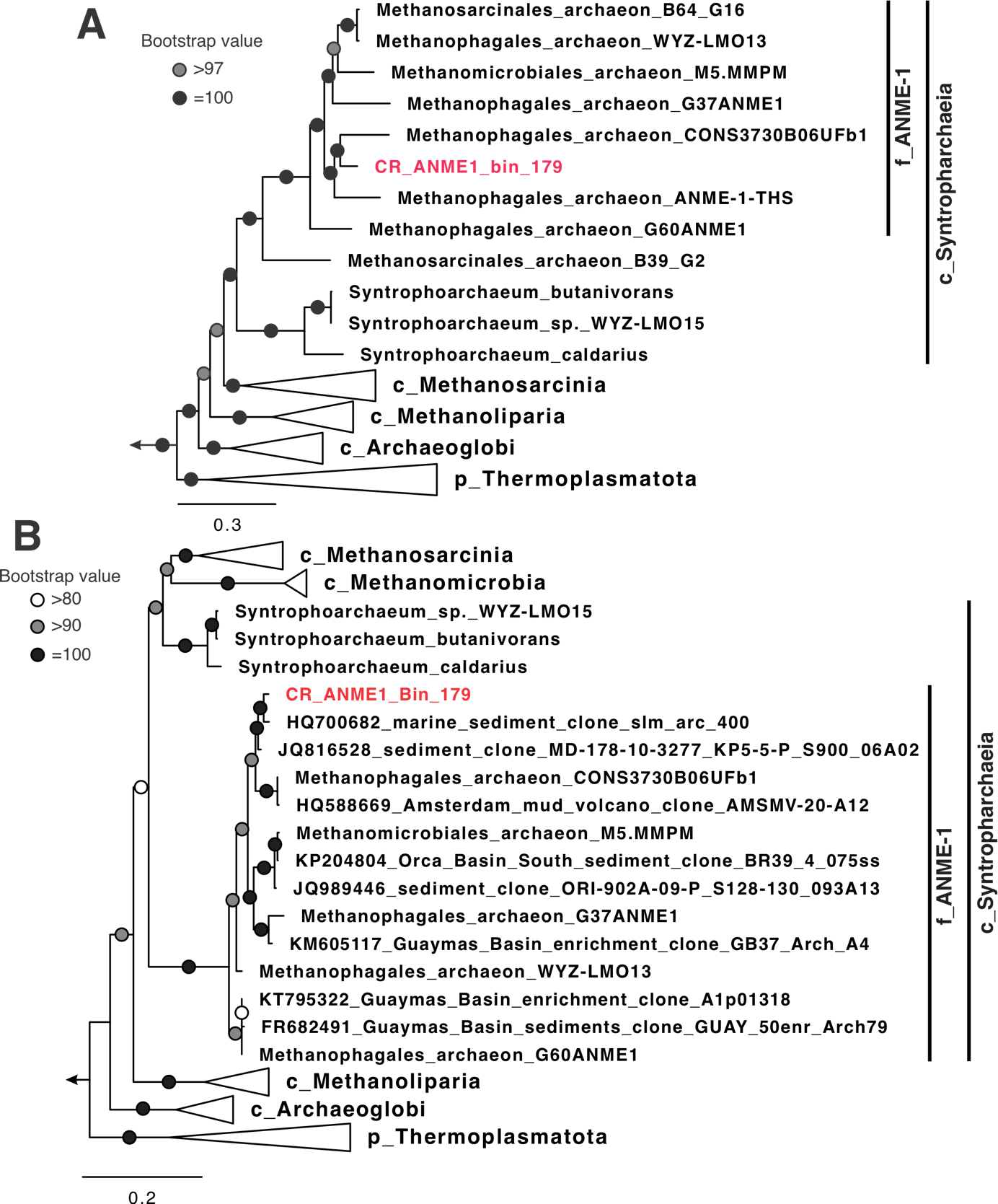

**Fig. S5. Maximum-likelihood phylogenetic tree of ANME-1 archaea based on 16 concatenated ribosomal proteins (A) and 16S rRNA gene (B).** **(A)** The trees were inferred using IQ-TREE v1.6.10 with the best-fit evolutionary models (i.e., the LG+R7 evolutionary model for **(A)** and SYM+R6 for **(B))** and 1000 ultrafast bootstraps. In both trees, the ANME-1 genome recovered in this study (CR_ANME1_Bin_179) is highlighted in red. The phylogenetic breadths of the class of Syntropharchaeia and the family of ANME-1 proposed by GTDB (https://gtdb.ecogenomic.org/) are also shown. The scale bars show estimated substitutions per residue.

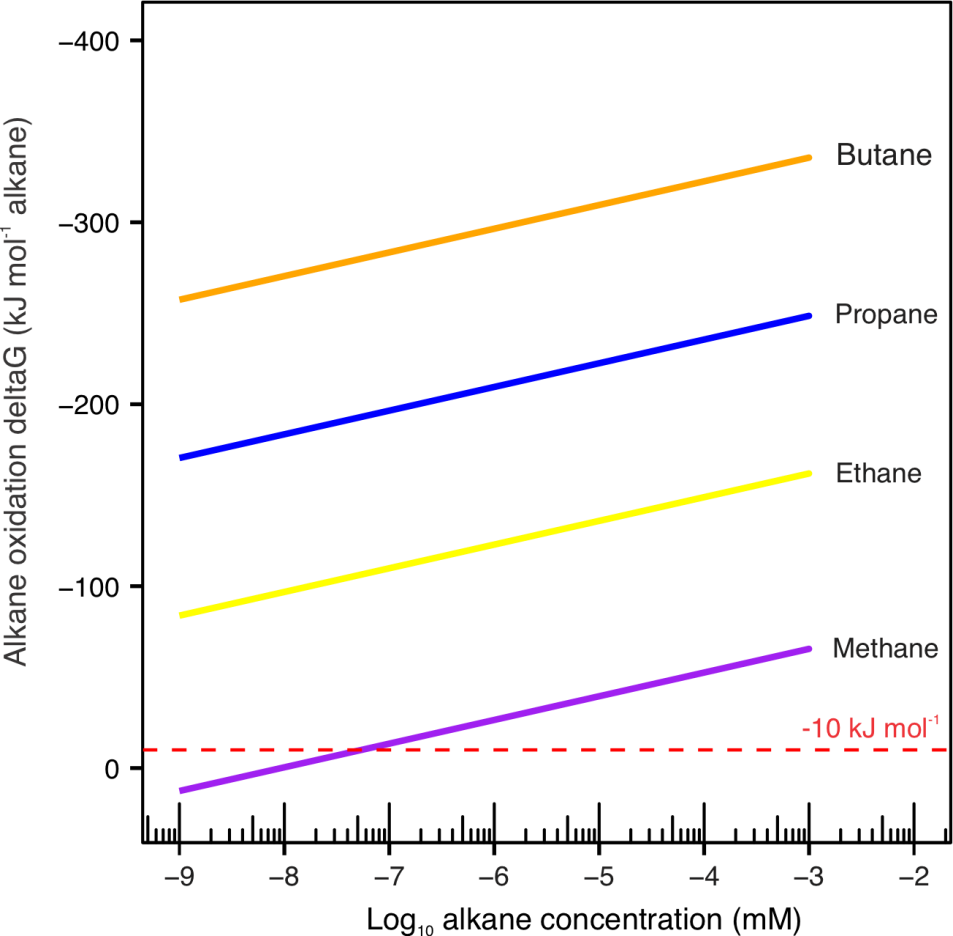

**Fig. S6. Gibbs free energy of alkane oxidation coupled to sulfate reduction.** The chemical equations of the reactions were shown in Table S3. A wide range of alkane concentrations (10^-9^ – 10^-3^ mM) were consider for each reaction, while the concentrations of the other reactants were kept constant (see the Materials and Method section for details).

**Table S1. Average nucleotide identity (ANI) and average amino acid identity (AAI) between genomes in the Candidatus Helarchaeota phylum**

|  | Hel_GB_A | Hel_GB_B | CR_Bin_097 | CR_Bin_143 | CR_Bin_291 |
| --- | --- | --- | --- | --- | --- |
| **ANI** |  |  |  |  |  |
| Hel_GB_B | <70% |  |  |  |  |
| CR_Bin_097 | 75% | <70% |  |  |  |
| CR_Bin_143 | <70% | 75% | <70% |  |  |
| CR_Bin_291 | 76% | <70% | 80% | <70% |  |
| SZ_4_Bin10_384 | <70% | 75% | <70% | 75% | <70% |
| **AAI** |  |  |  |  |  |
| Hel_GB_B | 53% |  |  |  |  |
| CR_Bin_097 | 69% | 51% |  |  |  |
| CR_Bin_143 | 53% | 51% | 54% |  |  |
| CR_Bin_291 | 70% | 51% | 75% | 54% |  |
| SZ_4_Bin10_384 | 52% | 51% | 52% | 54% | 52% |

**Table S2 Average nucleotide identity (ANI) and average amino acid identity (AAI) between genomes in the family of ANME1**

|  | ANME1_bin_179 | M5.MMPM | ANME-1-THS | CONS3730B06UFb1 | G60ANME1 | WYZ-LMO13 |
| --- | --- | --- | --- | --- | --- | --- |
| **ANI** |  |  |  |  |  |  |
| M5.MMPM | 79% |  |  |  |  |  |
| ANME-1-THS | 77% | <70% |  |  |  |  |
| CONS3730B06UFb1 | 78% | 78% | 76% |  |  |  |
| G60ANME1 | 77% | 78% | <70% | 77% |  |  |
| WYZ-LMO13 | 79% | 85% | 76% | 78% | 83% |  |
| B64_G16 | 79% | 80% | <70% | 77% | 79% | 88% |
| **AAI** |  |  |  |  |  |  |
| M5.MMPM | 75% |  |  |  |  |  |
| ANME-1-THS | 71% | 68% |  |  |  |  |
| CONS3730B06UFb1 | 72% | 72% | 68% |  |  |  |
| G60ANME1 | 69% | 69% | 66% | 67% |  |  |
| WYZ-LMO13 | 75% | 83% | 69% | 71% | 73% |  |
| B64_G16 | 76% | 77% | 69% | 71% | 70% | 87% |

**Table S3. Reactions of sulfate-dependent alkane oxidation included in**

**the thermodynamic calculation**

| **Reactions** | **Chemical equations** | **References** |
| --- | --- | --- |
| Ethane oxidation | 4[C_2_H_6_]+7[SO_4_^2-^]+6[H^+^] -> 8[HCO_3_^-^]+7[H_2_S]+4[H_2_O] | [21] |
| Propane oxidation | 2[C_3_H_8_]+5[SO_4_^2-^]+4[H^+^] -> 6[HCO_3_^-^]+5[H_2_S]+2[H_2_O] | [22] |
| Butane oxidation | 4[C_4_H_10_]+13[SO_4_^2-^]+10[H^+^] -> 16[HCO_3_^-^]+13[H_2_S]+4[H_2_O] | [23] |
| Methane oxidation | [CH_4_]+[SO_4_^2-^]+[H^+^] ->[HCO_3_^-^]+[H_2_S]+[H_2_O] | [24] |
